# Detection of Nontoxic BoNT/A Levels in Post-Facial Botox Injection Breastmilk using a Multi-technique Approach

**DOI:** 10.1101/2024.05.22.595434

**Authors:** Helene Gu, Zhenyu Xu, Renata Koviazina, Pengcheng Tan, Changcheng Zheng, Ferdinand Kappes, Domna G. Kotsifaki, Fangrong Shen, Anastasia Tsigkou

**Affiliations:** Reproductive Oncology Lab, Division of Natural and Applied Sciences, Duke Kunshan University, 8 Duke Ave, Kunshan, Jiangsu Province, 215316, China; Photonics Lab, Division of Natural and Applied Sciences, Duke Kunshan University, 8 Duke Ave, Kunshan, Jiangsu Province, 215316, China; Division of Natural and Applied Sciences, Duke Kunshan University, 8 Duke Ave, Kunshan, Jiangsu Province, 215316, China; Optical Characterization Lab, Division of Natural and Applied Sciences, Duke Kunshan University, 8 Duke Ave, Kunshan, Jiangsu Province, 215316, China; First Affiliated Hospital, Soochow University Department of Obstetrics and Gynecology, 296 Shizi St, Cang Lang Qu, Suzhou, Jiangsu Province, 215005, China

**Keywords:** Breastmilk, Breastfeeding, Botulinum Neurotoxin Type A, Botox, BoNT/A, ELISA, Western Blot, confocal micro-Raman Spectroscopy, Mass Spectrometry (LC-MC)

## Abstract

**Background:** The use of cosmetic Botox (Botulinum Neurotoxin Type A, BoNT/A ) has become increasingly prevalent. Particularly after pregnancy, postpartum depression represents one major factor motivating women to use Botox even during the lactation and breastfeeding period. Currently, there is limited understanding of the impact of Botox on lactation and the potential of its active component passing into breastmilk and affecting the infant.

**Methods:** Breastmilk samples were acquired from five women aged between 28 - 45 through a clinic in Suzhou, Jiangsu, P.R. China. Three sample sets ranged from 1 hour to 1 year after facial Botox treatments (64 U), whereas the remaining two sample sets were from women who never received Botox treatment. BoNT/A concentrations in samples were detected using standard Enzyme-Linked Immunosorbent Assay (ELISA), unreduced and reduced Western Blotting, confocal micro-Raman Spectroscopy, and Mass Spectrometry(LC-MS).

**Findings:** From ELISA, breastmilk BoNT/A concentrations peaked at 33.4 pg/mL 4 days after Botox injection. BoNT/A concentrations were highest overall in the first week and around two months after injection. While non-reducing polyacrylamide gel electrophoresis (PAGE) showed a protein band of 150 kDa peaking at 48 hours, reduced SDS-PAGE detected a 100 kDa protein first peaking at 72 hours, then re-emerging after 7 days, respectively, and in line with previous observations by others. Interestingly, micro-Raman spectroscopy indicated additional Raman peaks at 6 hours and 48 hours that were not present in other breastmilk samples which were evaluated in this study. However, no clear indication of BoNT/A was detected in Mass Spectrometry (LC-MS).

**Interpretation:** The amount of BoNT/A in breastmilk peaks around 48 hours, and at 2 months after facial injection. Even over a year after injection, BoNT/A can be detected. However, all quantities of BoNT/A detected in this study are highly likely to be safe for infants. Additionally, our study suggests that alternative methods, besides ELISA, may be utilized for the rapid detection of low concentrations of BoNT/A in body fluid samples.

**Funding:** Duke Kunshan University Start-Up funds, Duke Kunshan University Undergraduate Studies Signature Work Research Grant, Synear and Wang-Cai Biochemistry grants, and Kunshan Municipal Government research funding.

**Graphical Abstract:** 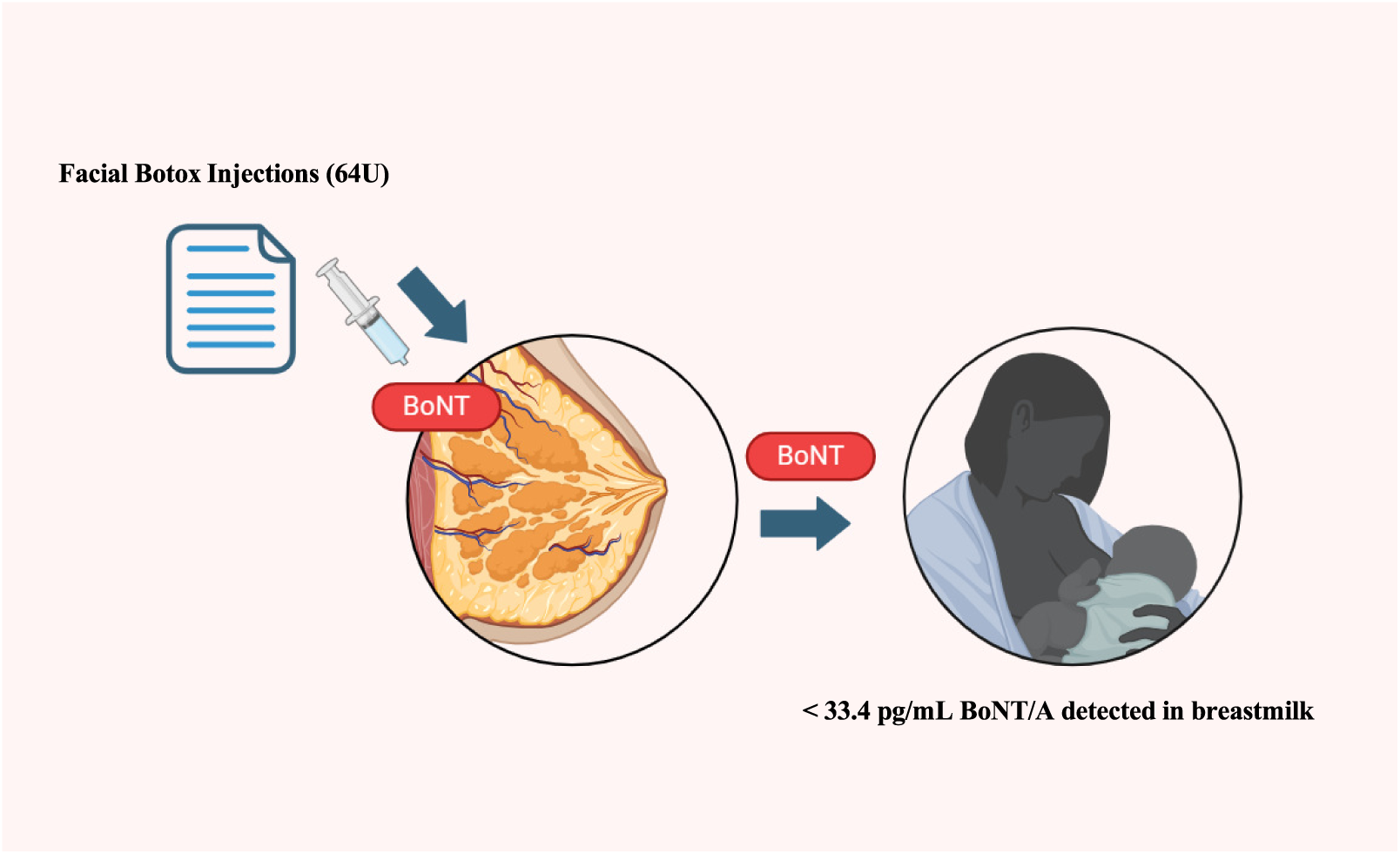

**Highlights:**

- BoNT/A was detected in the breastmilk of 3 women after facial Botox injections of 64 U.
- Toxin levels peaked in the first week and at around 2 months after injection.
- All detected levels (up to 33.40 pg/mL) were significantly below the lethal dose for newborns.

## 1. Introduction

In today’s social media oriented culture, caring about one’s appearance is becoming the new normal. Consequently, there has been a rapid growth of cosmetic procedures, with Botox, the most popular and easily accessible treatment, having developed into a seven billion dollar industry [1]. These nascent beauty conscious behavioral patterns also affect pregnant women, given that numerous studies have identified maternal depression in postnatal women [2–4]. During pregnancy, women’s hormones fluctuate and their bodies change with stomachs enlarging, breasts growing, ankles/wrists swelling, and skin gaining stretch marks as they grow a child. After pregnancy, the immense pressure to “bounce back” evokes body dissatisfaction and postnatal depression in roughly 10% of all women [2–4]. Without strong social support, low self-esteem can be a significant motivator for seeking timely Botox treatments, even during the early lactation period, which may represent a risk to the newborn.

Breastmilk, produced in the alveoli of breasts, derives its nutrients from the circulatory system, thus being prone to contamination by contents of the mother’s bloodstream. Despite this, the benefits of breastmilk are wide-spanning and well-studied. Breastfeeding benefits both the mother and newborn by reducing obesity, diabetes, hypertension, cardiovascular disease, and certain cancers [5, 6]. For the newborn, breastmilk from the first two days after birth provides high amounts of antibodies and white blood cells. Afterwards, it contains important proteins, sugars, vitamins, minerals, hormones, and growth factors [7, 8]. For the mother, breastfeeding can reduce stress, fatigue, and depression [9]. Therefore, the World Health Organization (WHO) recommends mothers initiating breastfeeding within the first hours of life and exclusively breastfeed for at least six months [10, 11]. However, exceptions are represented by cases of mothers with certain diseases or maternal medications proven to pose risks to babies [12].

In clinical treatments, Botox doses vary from 1.25 to 300 U [13, 14]. One U, *i.e* mouse unit, refers to the lethal dose that kills 50% of mice [15]. For humans, a lethal dose of Botox is around 3000 U [15]. Therefore, Botox injections have almost unanimously been found to be safe and free from long lasting side effects in clinical trials [13].

Conjectures have been made that BoNT/A, the active ingredient of Botox, is unlikely to enter breastmilk. Due to having a high molecular weight and high neurospecificity to its target, it is not likely to spread systemically [16]. However, many other drugs including antidepressants, cessation aids, and immune disease medications are detected in breastmilk [17, 18]. Some, such as Naltrexone, an alcohol/opioid substitute medication, are injected like Botox [17].

In the LactMed database, one recent study was conducted on Botox in four patients receiving facial injections. In two of the patients (receiving 54 U and 92 U, respectively), no BoNT/A was detected in breastmilk [19]. In the other two women, between 85.24 and 746.82 pg/mL were detected 0 - 5 days post-injection [19]. Other case reports related to Botulinum toxin and breastmilk are sparse and are related to maternal botulism from food, not Botox injections [17]. These cases are few in number, sample size, and time points measured.

Botox inhibits acetylcholine secretion in neurons of the peripheral nervous system, entering the nerve terminal almost immediately after injection [20–22]. The binding and internalization of Botox have a half-time of 10 minutes at body temperature, *i.e* 36*^◦^*C [23]. Within 12 hours after injection, it is either absorbed by nerve terminals or released from the muscles [8, 24]. The half-life for Botox in the circulatory system is several days before it reaches target organs [25]. Due to its high neurospecificity, it undergoes little biotransformation and does not accumulate in circulating cells, remaining largely in the free state [17]. The effects of Botox injection decrease after 2 months when it is fully metabolized, and it is generally at 3 - 4 months when effects completely wear off and a new injection is needed [13].

Given that an oral dose as small as 1 *µ*g/kg may be lethal, a lethal dose of Botox for a 3.5 kg baby would be 3.5 *µ*g [26]. In addition to lower body weight, newborns’ less developed immune systems may also make botulinum neurotoxin particularly dangerous, causing diminished facial expressions, loss of muscle tone, difficulty feeding, and general weakness [20]. However, if a recovery is made, there are generally no long-term effects [27].

Given the limited amount of studies, the Food Drug Administration (FDA) currently warns against pregnant and breastfeeding women using Botox [28]. Our study aims to inform this recommendation by quantifying BoNT/A levels in breastmilk post-injection via standard, *i.e* ELISA, Western Blotting and mass spectroscopy, and non-standard *i.e.* confocal micro-Raman spectroscopy, techniques.

## 2. Methods

### 2.1. Study Design and Participants

The study participants were three lactating women between the ages of 35 and 45 who underwent their most recent Botox treatment more than a year ago. Additionally, two breastfeeding women aged 28 to 32 who have never received Botox treatment served as controls (Table 1). Women received BoNT/A injections (BOTOX® Cosmetic onabotulinumtoxinA) treatment for glabellar lines, lateral canthal lines, and forehead lines in accordance with the FDA Botox medication guide [29]. They received 20 U, 24 U, and 20 U at the 3 sites respectively, totaling to 64 U per woman. All women were instructed to express 200 mL of breastmilk into separate labeled sterile bags (MEDELA) before treatment and 1, 6, 24, 36, 48, and 72 hours after treatment using a manual pump. Samples were immediately frozen in participants’ home freezers at *−*20*^◦^*C and transported to the laboratory at day 4 using dry ice and were stored *−*80*^◦^*C until analysis. In addition, further samples were collected at 5, 6, 7, 10, and 11 days, 2 months, 4 months, and 5 months following the same above procedure, under which botulinum neurotoxin can be stable for years [30].

**Table 1:** Summary of participant data and samples collected from 3 women who have received facial Botox injections and 2 women who have never received Botox.

| Woman | Injection Type | Age | Breastmilk Samples collected | Volume collected |
| --- | --- | --- | --- | --- |
| 1 | Facial (64 U) | 35 - 45 | 1 h, 6 h, 24 h, 36 h, 48 h, 72 h, 4 d, 5 d, 6 d, 11 d, 2 m, 4 m, 1+year | 200 mL each |
| 2 | Facial (64 U) | 35 - 45 | 6 h, 24 h, 36 h, 48 h, 72 h, 5 d, 7 d, 1+year | 200 mL each |
| 3 | Facial (64 U) | 35 - 45 | 24 h, 36 h, 48 h, 4 d, 5 d, 6 d, 11 d, 1+year | 200 mL each |
| 4 | Never received Botox | 28 | Normal breastmilk random sample | 100 mL |
| 5 | Never received Botox | 32 | Normal breastmilk random sample | 100 mL |

### 2.2. Sample Preparation and Dialysis

Breastmilk sample aliquots were centrifuged at 1,500 *×* g at 4*^◦^*C for 15 min and the skim milk fraction was collected through a disposable 5 mL needle syringe. The skim milk sample was centrifuged again at 3,000 *×* g at 4*^◦^*C for 30 min to remove remaining fat globules and cell debris. Samples were stored as 1 mL aliquots at *−*80*^◦^*C for further analysis.

Dialysis of breastmilk was performed overnight at 4*^◦^*C in a cold room with 50 kDa MWCO dialysis membranes (Shyuanye) and deionized (DI) water as the solute. Anti-BoNT/A (AbCam) antibodies were bound and then covalently cross-linked to the bead-immobilized protein A (Beyotime Protein A&G)). To produce a stable immunomatrix, the Seize X Immunoprecipitation Kit (Pierce, product #45215) was used as described by the manufacturer. The beads were centrifuged and washed three times with 500 *µ*L “Gentle Elution buffer” and two times with 500 *µ*L “Gentle Binding” buffer (Pierce, product #21030). Spin Cup columns with functionalized beads were wrapped with parafilm and stored at 4*^◦^*C until used. Breastmilk proteins were subjected to affinity chromatography with immobilized antibodies and were rotated at 8 rpm at 4*^◦^*C overnight, then centrifuged at 1700 *×* g for 10 min, and washed three times with 500 *µ*L of PBS buffer.

### 2.3. Enzyme-Linked Immunosorbent Assay (ELISA)

Breastmilk BoNT/A concentrations were measured in duplicates by a commercial ELISA kit for Human Botulinum Toxin Type A (BTX-ELISA China). ELISA was performed on samples spanning from 6 hours to 5 months following the manufacturer’s instructions and as outlined (Table 2).

**Table 2:** Breastmilk samples loaded in ELISA plate wells. Each sample was loaded in duplicate. The first row consists of the standards provided by the manufacturer to produce the standard curve.

|  | 1 | 2 | 3 | 4 | 5 | 6 | 7 | 8 | 9 | 10 | 11 | 12 |
| --- | --- | --- | --- | --- | --- | --- | --- | --- | --- | --- | --- | --- |
| **A** | ST 0<br>(0 pg/mL) |  | ST 1<br>(5 pg/mL) |  | ST 2<br>(10 pg/mL) |  | ST 3<br>(20 pg/mL) |  | ST 4<br>(40 pg/mL) |  | ST 5<br>(50 pg/mL) |  |
| **B** | Woman 1<br>6 h |  | Woman 1<br>24 h |  | Woman 1<br>36 h |  | Woman 1<br>48 h |  | Woman 1<br>72 h |  | Woman 1<br>14 days |  |
| **C** | Woman 1<br>5 days |  | Woman 1<br>7 days |  | Woman 1<br>11 days |  | Woman 1<br>2 months |  | Woman 1<br>4 months |  | Woman 1<br>5 months |  |
| **D** | Woman 4<br>no Botox |  | Woman 5<br>no Botox |  | Woman 2<br>24 h |  | Woman 3<br>36 h |  | Woman 3<br>48 h |  | Woman 3<br>72 h |  |

#### 2.3.1. Statistical Analysis

The ELISA plate was read by a Varioskan LUX microplate reader (Thermo Fisher Scientific, Inc.). Values of each pair of duplicates were averaged. BoNT/A concentration versus absorbance of the standard curve was plotted following the manufacturer’s manual. The absorbance of standard 1 (0 pg/mL) was subtracted from all other absorbance values. From these adjusted average absorbances, concentrations of the breastmilk samples were determined.

### 2.4. Polyacrylamide Gel Electrophoresis (PAGE) and Western Blot

#### 2.4.1. Nonreducing PAGE

In the first experiment, samples from 24, 36, and 48 hours were subjected to polyacrylamide gel electrophoresis using non-reducing conditions. In this procedure, the toxin shows only the protein band formed by units of molecular weight at 150 kDa.

#### 2.4.2. Reducing PAGE

In the second experiment, samples across a longer span of time were subjected to reducing electrophoresis. In reducing PAGE, the toxin is resolved into its heavy (molecular weight = 100 kDa) and light (molecular weight = 50 kDa) subunits due to breakage of disulfide bonds [31].

#### 2.4.3. Western Blotting

For all Western Blots, total protein concentration was quantified using BCA Protein assay (Thermo Fisher Scientific, Inc.). A total of 30 *µ*g of proteins in breastmilk samples were separated on 8 or 10% Tris-Glycine gels (Titan Scientific). Before loading, proteins were denatured using 3 *µ*L 5*×* SDS (Sodium Dodecyl Sulfate) and heated at 95*^◦^*C for 5 mins. For reduced PAGE, 3 *µ*L of beta-mercaptoethanol was also added before heating in order to break the disulfide bridge connecting the heavy (100 kDa) and light (50 kDa) chains of BoNT/A. This was not added to the unreduced PAGE.

After (SDS) - electrophoresis, proteins were transferred onto a polyvinyldiene fluoride (PVDF) membrane in a Mini Trans-Blot cell (Bio-Rad, CA, USA) at a constant current of 315 mA on ice for 90 min. The membrane was incubated for one hour at room temperature with a blocking solution of PBS/BSA 5%+*N aN*_3_, then washed for 5 min in Tris-buffered saline (TBS: 150 mM NaCl, 10 mM Tris-HCl (pH 7.4)), to remove the excess BSA (Bovine Serum Albumin protein). After blocking, mouse anti-clostridium botulinum toxin A [B364M] (ab252737) primary antibody diluted in 2% BSA/PBS was added to the membrane and incubated overnight at 4*^◦^*C on a shaker. The membrane was then washed once with Tris-buffered saline-Tween (TBST: 150 (AbCam) mM NaCl, 10 mM Tris-HCl, and 0.1% Tween 20) for 15 min, followed by three washes with TBS (15 min each). The membrane was then incubated at room temperature for 1 hour with a clean blot secondary antibody (1 : 3000 in TBS) (Sigma-Aldrich Inc.) to suppress signals from antibodies naturally present in human breastmilk [32]. The membrane was then washed once in TBST for 15 min and 4 times in TBST for 5 min and left in AP buffer (100 mM Tris-HCl (pH 9.5), 100 mM NaCl, and 5 mM MgCl_2_) for 5 min. The bands were detected with a ChemiDoc Imaging system (Bio-Rad) and normalized to the expression of GADPH by semi-quantitative analysis.

Applicable controls including loading control, negative control lysate, and no primary antibody control were conducted in accordance with the antibody producer’s protocol.

### 2.5. Confocal micro-Raman Spectroscopy

Raman spectra were collected at room temperature using a confocal micro-Raman Spectrometer (LabRAM HR Evolution, HORIBA Scientific). A diode laser beam at 532 nm with power at the sample plane of 3.2 mW was used to provide better Raman efficiency compared to longer wavelengths. The laser beam was focused using a high numerical aperture (NA = 1.25) oil immersion objective lens (Plan N 100*×*, Olympus) onto the sample to maximise the Raman signal. The spectrometer has a spectral resolution of 1.0 *cm^−^*^1^ and a spectral range of 100 - 4000 *cm^−^*^1^. Each spectrum was averaged from three scans with an integration time of 15 s. Using adhesive microscope spacers with a thickness of 120 *µ*m, a microwell was formed on the microscope glass slide, and a spectrum occurred in the 6 *µ*L sample solution under a glass cover slide. The Raman spectra were analyzed using LabSpec6 software to identify the peaks. The analysis included baseline correction before peak position determination. Then the Savitzky–Golay filter was used for spectra smoothing by a 3*^rd^*-degree polynomial function.

### 2.6. Liquid Chromatography - Mass Spectrometry (LC-MS)

Liquid Chromatography - Mass Spectrometry (LC-MS) was used to analyze unprocessed breastmilk samples of 1 mL taken at 6 and 48 hours post-Botox administration. Liquid Chromatography was conducted using a Vanquish Neo UPLC system (Thermo Fisher Scientific, Inc.). The Mass spectrometry was analyzed using an Orbitrap Astral with a nano-electrospray ion source. Analysis methods including Gene Ontology (GO), Domain Annotation, KEGG, Subcellular Localization, COG/KOG, Reactome, WikiPathways, Hallmark Signature Gene sets, and Transcription Factor annotation to identify and determine the functions of proteins. Mass spectrometry was outsourced to the company PTM Bio Suzhou China.

## 3. Results

### 3.1. ELISA Results

Using a commercially available ELISA kit, concentrations of BoNT/A ranging from 0 to 33.40 pg/mL were detected in all breastmilk samples of women that received Botox treatment, while no BoNT/A was detected in the control group (Fig. 1). The highest concentration observed was in the sample from woman 1, 4 days after the injection (33.40 pg/mL). This is in line with the Hudson *et al.* observations that found peaks at 3 and 5 days, respectively, in two women [19]. We observed that after the first week, circulating BoNT/A concentrations decrease to a low concentration of 6.88 pg/mL at 11 days, before another peak of 26.55 pg/mL at 2 months (Fig. 1).

**Figure 1:**
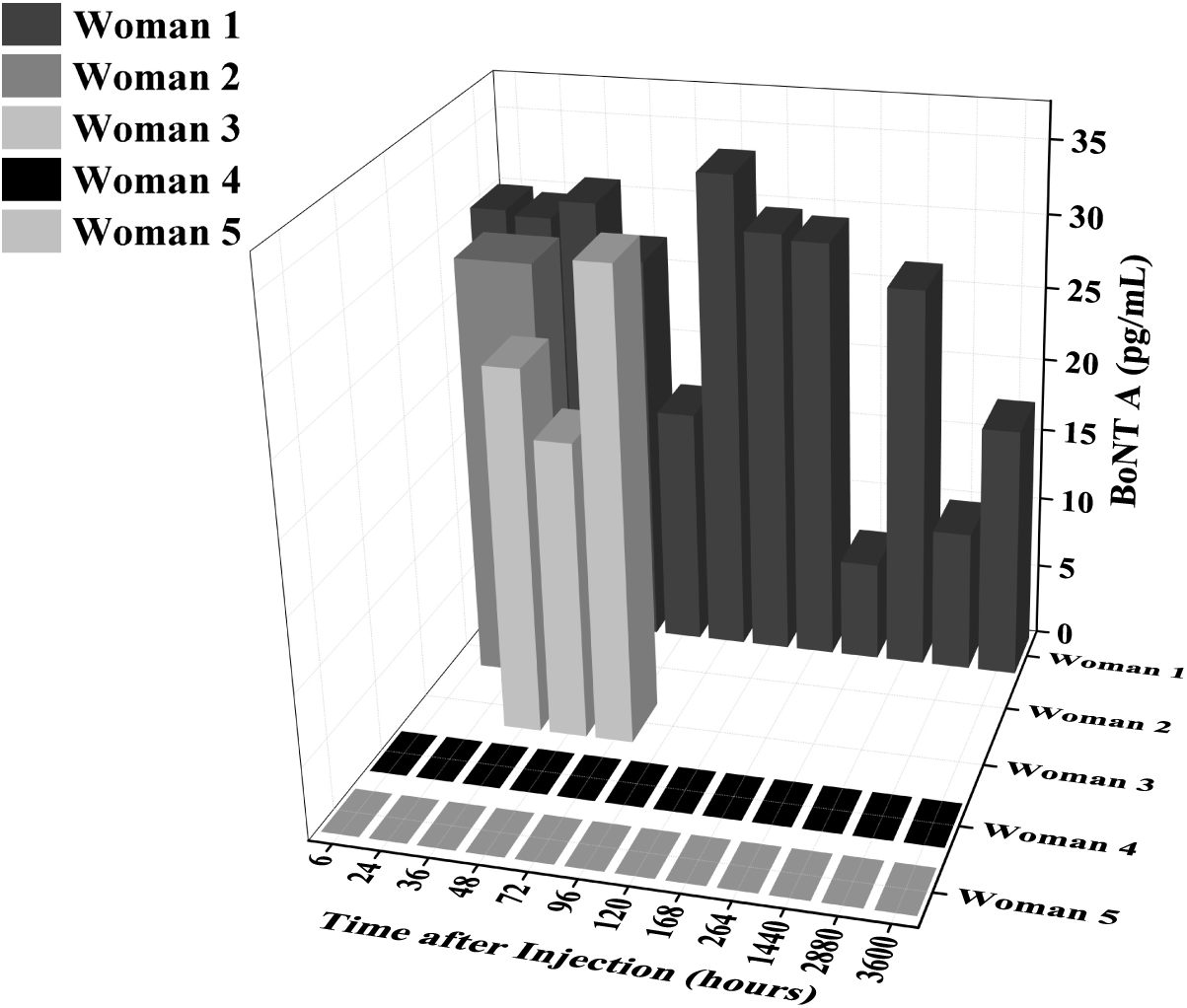
Concentrations of BoNT/A in ELISA samples as a function of time and as a function of breastmilk samples. The level of BoNT/A in breastmilk generally decreases over time after the first week (6 - 168 hours) but has a distinct peak again at 2 months (1440 hours).

### 3.2. Western Blot Results

Western blot analyses were conducted to apply an alternative method for detection of BoNT/A proteins in the breastmilk of the three lactating women.

For the loading control, clear bands corresponding to Glyceraldehyde-3-phosphate dehydrogenase (GAPDH) were observed at around 37 kDa when 10 *µ*L of breastmilk was analyzed, confirming that the protein concentration was sufficient and that breastmilk samples were adjusted to similar total protein content (Fig. 2(A). In the negative control lysate, as illustrated in the first wells of Figure 2 (C), samples from woman 4 and woman 5, who have never had received treatment of BoNT/A showed the absence of 150 kDa marker for BoNT/A.

**Figure 2:**
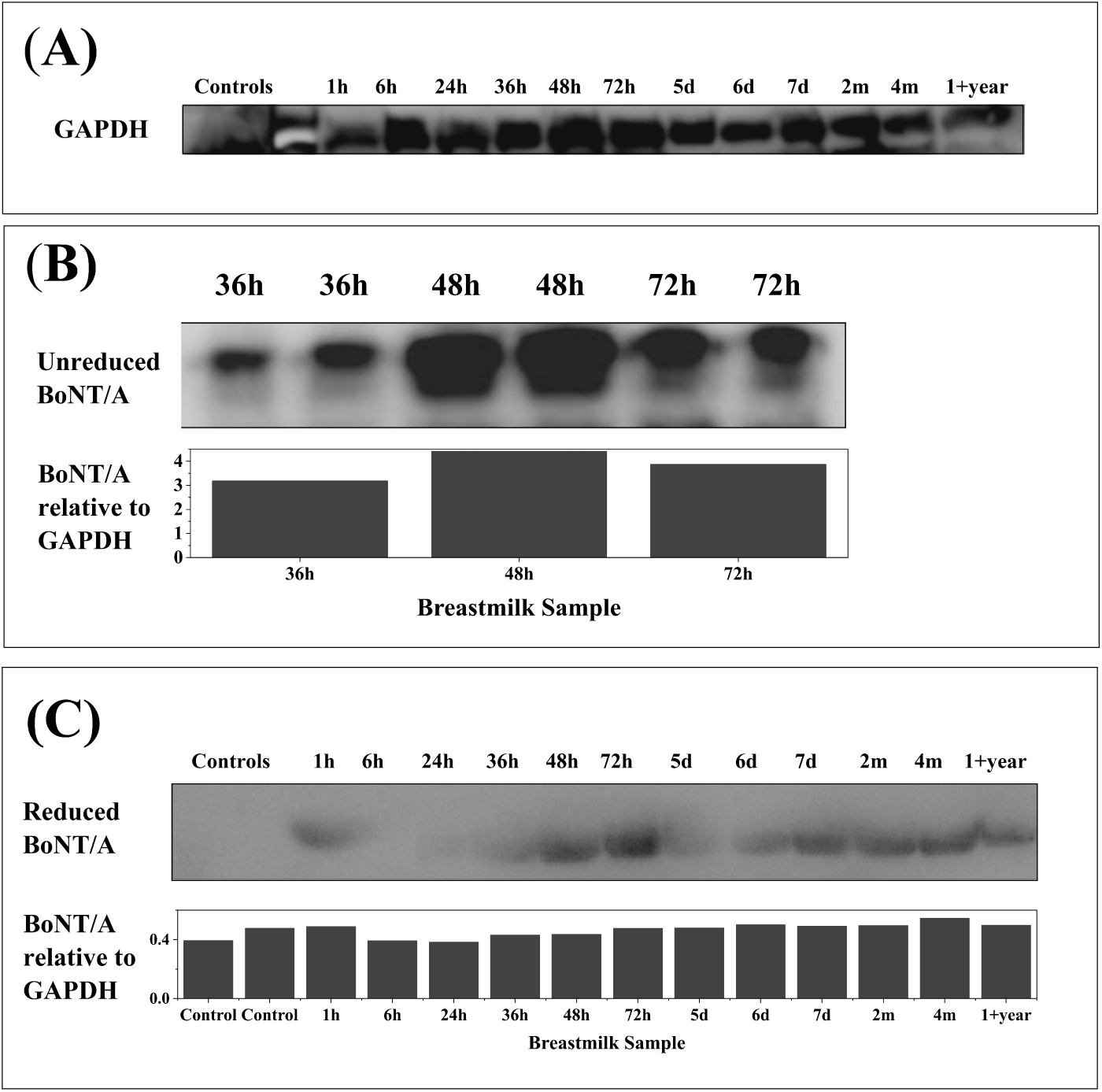
Western Blot Analysis of Breastmilk Reveals Presence of BoNT/A. **(A)** Glyceraldehyde-3-phosphate dehydrogenase (GADPH) loading control. GADPH was relatively constant in all samples, indicating comparable total protein concentrations of breastmilk. **(B)** Unreduced Western Blot analysis of woman 1’s breastmilk samples from 36, 48, and 72 hours. Bands around 180, 150, 120, and 45 kDa were observed. The band at around 150 kDa was most prevalent at 48 hours. Concentrated and purified woman 1 post Botox samples were run with 1 : 1000 Botulinum primary antibody, no beta mercaptoethanol, and 1 : 500 clean blot secondary antibody on a 10% gel. **(C)** Western blot analysis across a larger range of time from woman 1 was run again with an 8% gel under the same conditions. A band between at 100 kDa was observed. It was present the 1 hour, 36 hours, 48 hours, 72 hours, 5 days, 6 days, 7 days, 2 months, 4 months, and 1+ year samples but absent in the samples of women who had not received Botox before. It was also not present at 6 hours and 24 hours. This is the same molecular weight as the heavy chain of BoNT/A, between 85 - 100 kDa under reducing conditions [35]. Bands around 150 kDa and 45 kDa were also observed.

The Western blots revealed the presence of fluctuating 150 kDa (Fig. 2 (B)) and 100 kDa (Fig. 2 (C)). As BoNT/A protein has a molecular weight of 150 kDa, consisting of a light chain (50 kDa) and a heavy chain (100 kDa); both chains have different functions in the neurotoxin’s mechanism of action [33]. The detection utilized a primary antibody specific to the 1280 - 1292*^th^* amino acid residues of the heavy chain of the BoNT/A protein. To confirm the presence of sulfide bonds we compared between conducting reducing and non-reducing PAGE, anticipating the identification of a product approximately 150 kDa (non-reducing PAGE) or 100 kDa (reducing PAGE) in line with previous studies [34]. BoNT/A is produced as a single-chain polypeptide (150 kDa) that is inactive. Proteases nick the polypeptide chain resulting in a toxin that is pharmacologically active and consists of two chains: a heavy chain (100 kDa) and a light chain (50 kDa) connected together by a disulfide bond.

#### 3.2.1. Unreduced Western Blot

In unreduced samples of breastmilk 36, 48, and 72 hours after injection, the highest concentration of bands at 150 kDa occurred at 48 hours (Fig. 2(A)).

#### 3.2.2. Reduced Western Blot

Protein bands around 100 kDa are present as early as one hour post-treatment. Notably, over a longer course of time, this band peaked again on day 7 and was persistent up to over 1 year post-treatment (Fig. 2(B)).

### 3.3. Raman Spectroscopy Results

Next we were interested in investigating the BoNT/A concentration in breastmilk with a non-standard technique *i.e.*, confocal micro-Raman Spectroscopy. Figure 3 (A) and (B) show the Raman spectra of the samples collected from woman 1 and 2 (Table 1). Several characteristic peaks of breastmilk and BoNT/A were detected and summarized in Table 3. The presence of protein was identified by the amide I signature at 1656 cm*^−^*^1^ in relatively high intensity in sample 1 year compared with this band in 6 hours, 48 hours and 72 hours after Botox injection in case of the woman 1 and 2 (Fig. 3). Lipids are related to the presence of the hydrocarbon chain and can be observed in three regions: **(I)**:1200 - 1050 cm*^−^*^1^, **(II)**: 1300 - 1250 cm*^−^*^1^ and **(III)**: 1500 - 1400 cm*^−^*^1^ (Fig. 3) [36]. Specifically, the bands in the range of 1500 - 1400 cm*^−^*^1^ and 1300 cm*^−^*^1^ are linked to scissoring and twisting vibrations of the *CH*_2_ and *CH*_3_ groups, respectively, while the band in the region of 1200 - 1050 cm*^−^*^1^ are related to the *C −C* stretching vibrations [36, 37]. We observed that the peak around 1440 cm*^−^*^1^ that related to *δ*(CH_2_) scissoring vibrations [36, 37] show a high-intensity magnitude in both cases, *i.e.* woman 1 and woman 2, for the 1 year after Botox sample (Fig. 3(A)). In addition, the peak at 1156 cm*^−^*^1^ that is related to *δ*CH_2_ twisting vibrations [36, 37] has a relatively high-intensity magnitude in case of 48 hours and 6 hours after Botox injection samples for woman 1 and 2, respectively. Likewise, the peak at 1518 cm*^−^*^1^ which is related to *C − H* stretching vibrations [36, 37], shows a large intensity magnitude in the case of women 1 for 48 hours and 2 for 6 hours after Botox injection, in comparison with the corresponding one Raman peak observed on the rest of the breastmilk samples. Lastly, twisting and rocking vibrations of *CH*_2_ are noted in Raman spectrum at 870 cm*^−^*^1^ and 1156 cm*^−^*^1^ [37].

**Figure 3:**
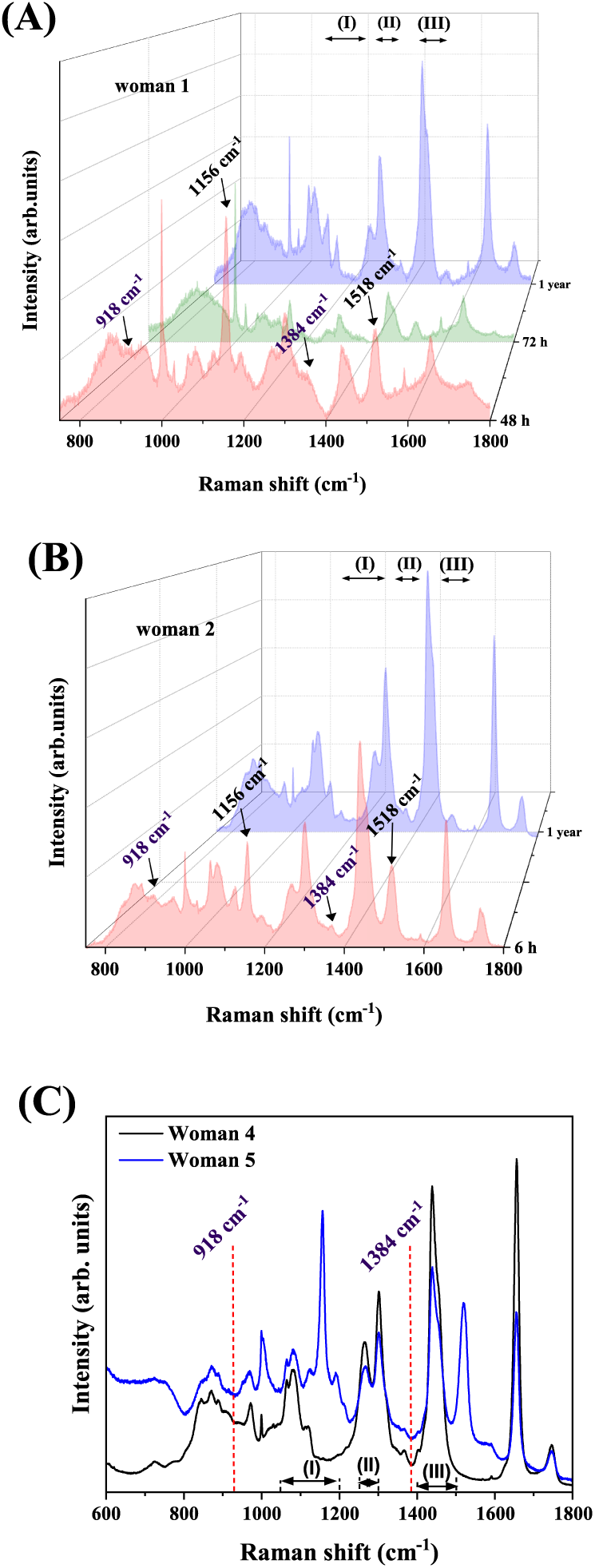
Raman Spectra of Breastmilk Samples from (A) woman 1, (B) woman 2 and (C) woman 4 and 5. The measurements were performed in a liquid environment with an excitation wavelength at 532 nm, an oil immersion objective lens with a magnification of 100*×*, and an integration time of 15 s. The characteristic peaks at 918 *cm^−^*^1^ and 1384 *cm^−^*^1^ were observed in samples 48 hours (woman 1) and 6 hours (woman 2) suggesting the presence of BoNT/A [38]. Note: **(I)**:1200 - 1050 cm*^−^*^1^, **(II)**: 1300 - 1250 cm*^−^*^1^ and **(III)**: 1500 - 1400 cm*^−^*^1^. The red dashed lines indicate the position of missing peaks in Raman spectrum of women 4 and 5.

**Figure 4:**
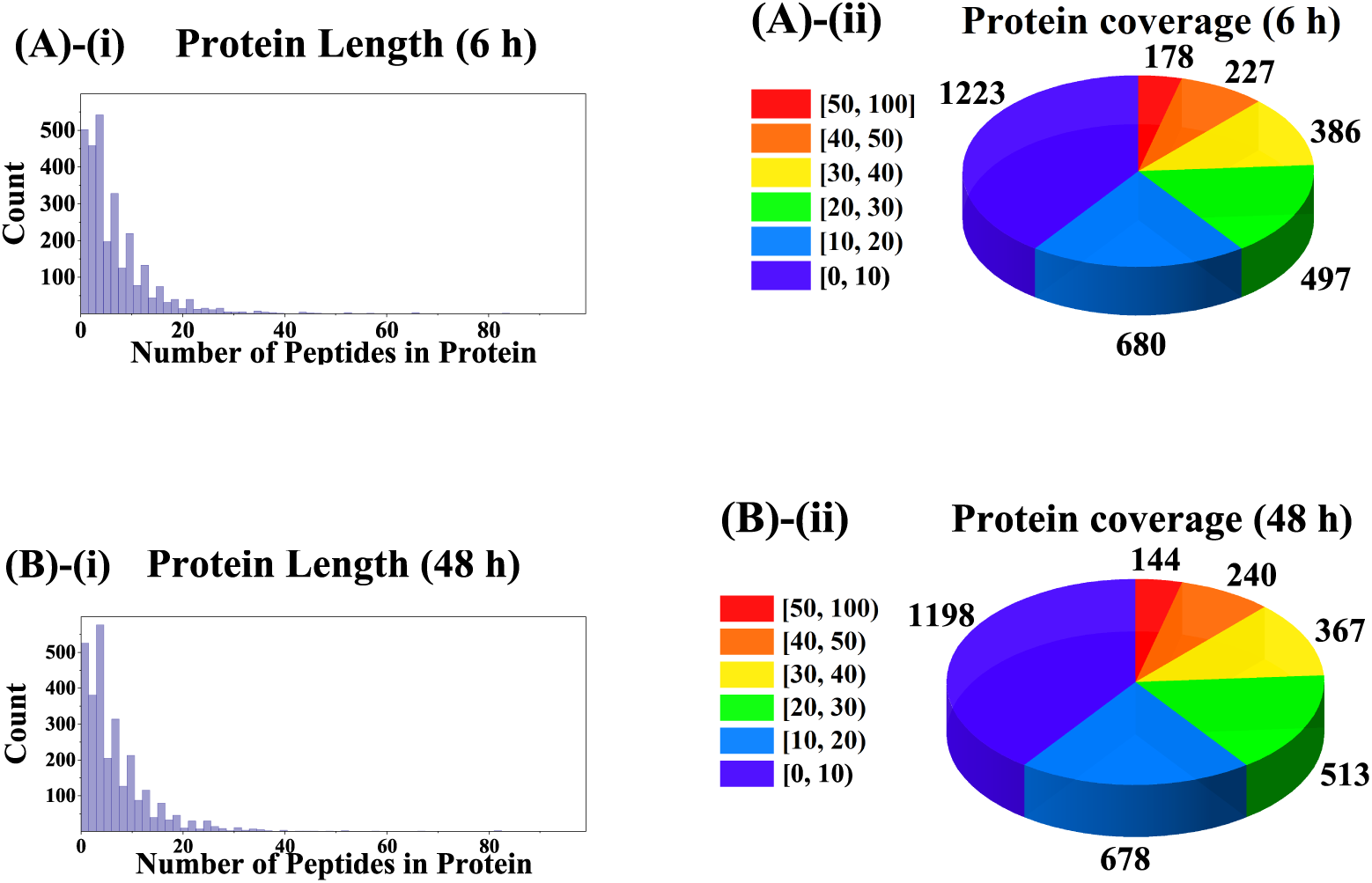
Peptide Length and Protein Coverage. Most proteins captured were less than 20 peptides long, and only a small portion of proteins were captured

**Table 3:** Raman Peaks of Breastmilk Samples [36, 37].

| Wavenumber ($cm^{-1}$) | Vibration |
| --- | --- |
| 850 - 920 | CH<sub>3</sub> rocking, CH<sub>2</sub> rocking<br>CC backbone, C-CH<sub>3</sub> stretching |
| 1000 | CC symmetric stretching |
| 1080 | CC asymmetric stretching |
| 1120 - 1123 | CC anti-sym stretching |
| 1437 - 1460 | $\delta(CH_2)$ scissoring, CH<sub>2</sub> deformation |
| 1656 | Amide I |
| 1749 | C=O stretching |

Comparing the spectra in Figures 3, we noted two Raman peaks around 918 *cm^−^*^1^ and 1384 *cm^−^*^1^ only in the samples of 48 hours (woman 1) and 6 hours (woman 2). These peaks have been detected in Clostridium Botulinum B bacterium which can produce Botox in specific conditions [38, 39].

### 3.4. Mass Spectrometry Results

In the 6 hours sample, 22666 peptides were detected, consisting of 22097 unique peptides and 3191 identified proteins. In the 48 hours sample, 22333 peptides were detected, with 21688 unique peptides and 3140 identified proteins. The majority of the protein fragments digested with Trypsin detected were less than 20 peptides. Therefore, most proteins were not fully covered (Fig. 4). BoNT/A was not detected among these peptides.

## 4. Discussion

Botox has been extensively used in clinical treatments for conditions such as excessive sweating (hyperhidrosis), muscle stiffness (spasticity), enlarged facial muscles (facial muscular hypertrophy), muscle spasms, migraines and cosmetic treatments [40]. As more women delay pregnancies and seek to continue breastfeeding beyond the WHO-recommended 2 years, and with others experiencing migraines or desiring post-pregnancy improvements, there is insufficient information available regarding the safety of using BoNT/A during breastfeeding for infants.

Here, we present four different methods for the detection and quantification of BoNT/A in breastmilk samples of three lactating women from 1 hour to over 1 year after Botox injection. These methods were a commercially available BoNT/A ELISA immunoassay kit, Western Blot analysis, Confocal micro-Raman Spectroscopy, and Mass Spectroscopy (LC-MS).

Almost all medications transfer from blood to breastmilk in varying amounts [17]. The Relative Infant Dose (RID), serves as the criterion for determining the safety of a medication’s concentration in breastmilk for an infant. It is important to note that concerns about the baby’s safety arise when the RID is greater than 10% [41]. The transfer of drugs into breastmilk is influenced by protein binding, lipid solubility, and ionization [42]. Heparin and Insulin were believed to be too large to cross biological membranes, but recent evidence shows otherwise [43, 44]. Likewise, the recent study from Hudson *et al.* found that 8 of the 16 samples had varying levels of botulinum toxin ranging from 85.24 to 746.82 pg/mL, while 8 samples had no toxin detected [19].

Pediatric doses of BoNT/A treatment for children with spastic Cerebral Palsy ranged from 12 to 16 U/kg and from 15 to 25 U/Kg of body weight for Botox and Dysport, respectively [45]. The effects of injection are seen at 3 to 6 days and the peak at 6 weeks [46]. When pregnant mice and rats were injected intramuscularly during the period of organogenesis, the developmental NOEL (No Observed Effect Level) of BoNT/A was 4 U/kg. Higher doses (*i.e.*, 8 or 16 U/kg) were associated with reductions in fetal body weights and/or delayed ossification [47].

Due to the current limited information on the detection of BoNT/A in breastmilk, we tested the above methods to provide a better sense of the risk versus benefit assessment.

LC-MS consists of 2 parts; Liquid chromatography (LC) is first used to separate components of a solution by mass, after which mass spectrometry (MS) is used to identify these components based on fragmentation pattern differences. It is highly sensitive, down to detection of parts per million (ppm), and can be performed quickly and efficiently [48]. However, in our results concentrations of BoNT/A were even lower than this threshold, requiring more targeted approaches.

Recent studies have shown that optical screenings and diagnoses have been widely used for characterizing biological media with reliable results including in differentiating between feeding male and female infants mother’s milk [49, 50]. By utilizing confocal micro-Raman spectroscopy, we collected and analysed the Raman spectrum of breastmilk samples in order to identify BoNT/A. Two Raman peaks around 918 *cm^−^*^1^ and 1384 *cm^−^*^1^ were noted only in the samples of 48 h (woman 1) and 6 h (woman 2). This observation along with the high-intensity magnitude of Raman peaks at 1156 *cm^−^*^1^ and 1518 *cm^−^*^1^ in our Raman spectra, lead us to assume the potential presence of BoNT/A in these two samples since these two peaks have been also detected in Clostridium botulinum B bacterium [38, 39]. However, additional experiments using more sensitive techniques such as Surface-Enhanced Raman Scattering (SERS) [51] or Fano-enhanced Raman scattering (FERS) [52] could provide complementary evidence to support this observation.

Western blot was performed to detect BoNT/A in breastmilk samples. The specificity of the band at molecular weight 150 kDa under no reducing PAGE and 100 kDa under reducing agent beta-mercaptoethanol. This was important to confirm the presence of sulfide bonds we compared between reducing and non-reducing PAGE bonds for BoNT/A was confirmed in all lactating women who received facial injections (woman 1, 2, 3), while no presence of BoNT/A was detected in women who never received Botox treatment (women 4,5). Circulating BoNT/A can be detected in the bloodstream even one year after Botox treatment, as the BoNT/A that had previously been rapidly uptaken by peripheral nerve terminals is gradually released back into the bloodstream. This is then excreted to the kidneys as waste or excreted into breastmilk [53], as we have shown in this study.

One unanticipated result was just how long BoNT/A can remain in the body. Due to prior treatment of the lactating women over 1 year before this study, the original negative “pre Botox” controls we used in Western blots were from women 1, 2, and 3. Because these women had all received Botox procedures for a time longer than one year ago, this led to inconclusive results and being unable to find differentiating molecular weight bands on the gel. However, upon acquiring breastmilk samples from women 4 and 5, who had never had Botox before in their lifetime, bands at 150 kDa were not observed. This indicates that even after over a year since a facial injection, detectable amounts of BoNT/A may still be present in circulation.

Enzyme linked immunoassays (ELISA) vary in sensitivity and specificity mainly due to different types of both detection and capture antibodies [54]. For instance, commercial ELISA kits tested different values of total corticosterone in the same serum samples [54]. With regards to the quantitative BoNT/A breastmilk concentrations determined by ELISA, our concentrations were significantly lower than the Hudson *et al.* study, by about 10-fold [19]. The findings observed may be connected to the higher doses of BoNT/A administered to the patients. Additionally, the variation in outcomes might be influenced by the ELISA kit which detects various epitopes. Metabolic variances in the breakdown of Botox could also have played a role. The body’s uptake, circulation, and excretion of BoNT/A is complicated, and metabolism can differ based on genetics or lifestyle choices. In particular, BoNT/A uptake to nerve endings increases with activity and temperature, reducing initial Botox concentrations but increasing later concentrations circulation[53].

Our study and previous ones have had a limited sample size and cover only facial Botox injection procedures. Future studies can be done to further elucidate how Botox metabolism changes with different injections and different lifestyles. However, given that the highest value was 33.40 pg/mL, this is 1 millionth of the lethal dose 3.5 *µ*g. Hence, for lethal concentrations, 1000 L would be required. Based on an average intake of 670 mL of breastmilk/day, a breastfed infant would ingest 0.50 *µ*g of Botox toxin in 24 hours. Therefore, it is unlikely that the risks of Botox levels in breastmilk are high enough to outweigh the benefits of breastmilk for the infant and mother.

Based on our knowledge, this study is the first to examine breastmilk from lactating women for presence of BoNT/A using Western Blot, confocal micro-Raman Spectroscopy, and Mass Spectrometry. Its results from ELISA are consistent with those of a recent publication using ELISA [19]. Together with the results from Hudson *et al.*, this study supports the conclusion that facial BoNT/A injections does not require the interruption of breastfeeding[19].

## 5. Authors Contribution

AT conceived the idea of this work, DGK conceived the idea and the experiments of Raman spectroscopy. AT and DGK supervised all stages of this work, oversaw the methodology, and conducted review and editing of the manuscript. HG, ZX, RK, and PT performed the experiments and analyzed the data. HG wrote the first draft. FK and CZ discussed the experimental data. All the authors contributed to writing the paper.

## Acknowledgements

The authors would like to thank Holly McClure for fruitful discussions and assisting with the Western Blot process.

## References

[1] W. W. Hood, C. S. Wilson, The literature of bibliometrics, scientometrics, and informetrics, Scientometrics 52 (2001) 291–314.

[2] N. I. Gavin, B. N. Gaynes, K. N. Lohr, S. Meltzer-Brody, G. Gartlehner, T. Swinson, Perinatal depression: A systematic review of prevalence and incidence, Obstetrics & Gynecology 106 (5) (2005) 1071–1083.

[3] A. C. Sweeney, R. Fingerhut, Examining relationships between body dissatisfaction, maladaptive perfectionism, and postpartum depression symptoms, Journal of Obstetric, Gynecologic & Neonatal Nursing 42 (5) (2013) 551–561.

[4] F. J. Riesco-González, I. Antuńez-Calvente, J. M. Vázquez-Lara, L. Rodríguez-Díaz, R. Palomo-Gómez, J. Gómez-Salgado, J. J. García-Iglesias, T. Parrón-Carrenõ, F. J. Fernández-Carrasco, Body image dissatisfaction as a risk factor for postpartum depression, Medicina 58 (6) (2022) 752.

[5] C. Binns, M. Lee, W. Y. Low, The long-term public health benefits of breastfeeding, Asia Pacific Journal of Public Health 28 (1) (2016) 7–14.

[6] P. A. R. Neves, F. S. Juliana S. Vaz, P. Baker, G. Gatica-Domínguez, E. Piwoz, N. Rollins, C. G. Victora, Rates and time trends in the consumption of breastmilk, formula, and animal milk by children younger than 2 years from 2000 to 2019: analysis of 113 countries, The Lancet Child and Adolescent Health (2021).

[7] F. Bhora, B. Dunkin, H. Aly, B. Bass, A. Sidawy, J. Harmon, Effect of growth factors on cell proliferation and epithelialization in human skin, Journal of Surgical Research 59 (2) (1995) 236–244.

[8] L. Ahmed, S. N. Islam, M. Khan, S. Huque, M. Ahsan, Antioxidant micronutrient profile (vitamin E, C, A, Copper, Zinc, Iiron) of colostrum: association with maternal characteristics, The Journal of Tropical Pediatrics and Environmental Child Health 50 (2004) 357–358.

[9] B. Figueiredo, C. Canário, I. Tendais, T. M. Pinto, D. A. Kenny, T. Field, Breastfeeding is negatively affected by prenatal depression and reduces postpartum depression, Psychological Medicine 44 (5) (2014) 927–936.

[10] Z. Tucker, C. O’Malley, Mental health benefits of breastfeeding: A literature review, Cureus 14 (9) (2022) e29199.

[11] World Health Organization (WHO), Infant and young child feeding, https://www.who.int/news-room/fact-sheets/detail/infant-and-young-child-feeding (2023).

[12] Centers for Disease Control and Prevention (CDC), When breastfeeding or feeding expressed milk is not recommended, https://www.cdc.gov/breastfeeding/breastfeeding-special-cir cumstances/ contraindications-to-breastfeeding.html (2023).

[13] F. Brandt, N. Swanson, L. Baumann, B. Huber, Randomized, placebocontrolled study of a new botulinum toxin type a for treatment of glabellar lines: efficacy and safety, Dermatologic Surgery 35 (12) (2009) 1893–1901.

[14] J. Morgan, S. Iyer, E. Moser, C. Singer, K. Sethi, Botulinum toxin A during pregnancy: A survey of treating physicians, Journal of Neurology, Neurosurgery & Psychiatry 77 (1) (2006) 117–119.

[15] P. T. Ting, A. Freiman, The story of Clostridium botulinum: from food poisoning to Botox, Clinical Medicine journal - London 4 (3) (2004) 258–261.

[16] G. Hilderbrand, C. Lamanna, R. Heckly, Distribution and particle size of type A Botulinum toxin in body fluids of intravenously injected rabbits, Proceedings of the Society for Experimental Biology and Medicine 107 (2) (1961) 284–289.

[17] H. C. Sachs, C. O. Drugs, The transfer of drugs and therapeutics into human breast milk: an update on selected topics, Pediatrics 132 (3) (2013) e796–809.

[18] M. Julsgaard, U. Mahadevan, T. Vestergaard, R. Mols, M. Ferrante, P. Augustijns, Tofacitinib concentrations in plasma and breastmilk of a lactating woman with ulcerative colitis, The Lancet Gastroenterology & Hepatology (2023).

[19] C. Hudson, P. Wilson, D. Lieberman, H. Mittelman, S. Parikh, Analysis of breast milk samples in lactating women after undergoing Botulinum toxin injections for facial rejuvenation: A pilot study, Facial Plastic Surgery & Aesthetic Medicine (2024).

[20] A. M. O. Bakheit, C. D. Ward, D. L. McLellan, Generalised botulism-like syndrome after intramuscular injections of Botulinum toxin Type A: a report of two cases, Journal of neurology, neurosurgery, and psychiatry 62 (2) (1997) 198.

[21] A. Carruthers, J. Carruthers, S. Said, Dose-ranging study of Botulinum toxin type A in the treatment of glabellar rhytids in females, Dermatologic Surgery 31 (4) (2005) 414–422.

[22] E. J. Schantz, E. A. Johnson, Botulinum toxin: the story of its development for the treatment of human disease, Perspectives in Biology and Medicine 40 (3) (1997) 317–327.

[23] J. D. Black, J. O. Dolly, Interaction of 125I-labeled Botulinum neurotoxins with nerve terminals. II. autoradiographic evidence for its uptake into motor nerves by acceptor-mediated endocytosis, Journal of cell biology 103 (2) (1986) 535–544.

[24] V. B. Brooks, The action of Botulinum toxin on motor-nerve filaments, The Journal of Physiology 123 (3) (1954) 501.

[25] F. H. Al-Saleem, D. M. Ancharski, E. Ravichandran, S. G. Joshi, A. K. Singh, Y. Gong, L. L. Simpson, The role of systemic handling in the pathophysiologic actions of Botulinum toxin, Experimental Therapeutics 326 (3) (2008) 856–863.

[26] S. S. Arnon, R. Schechter, T. V. Inglesby, D. A. Henderson, J. G. Barlett, M. S. Ascher, E. Eitzen, A. D. Fine, J. Hauer, M. Layton, S. Lillibridge, M. T. Osterholm, T. O’Toole, G. Parker, T. M. Perl, P. K. Russell, D. L. Swerdlow, K. Tonat, Botulinum toxin as a biological weapon: medical and public health management, JAMA 285 (8) (2001) 1059–1070.

[27] I. Brook, Infant Botulism, Journal of perinatology 27 (3) (2007) 175–180.

[28] Medication Guide BOTOX Cosmetic for Injection (2011).

[29] BOTOX (onabotulinumtoxinA) for injection, for intramuscular, intradetrusor, or intradermal use, accessed on April 14, 2024 (1989).

[30] C. Malizio, M. Goodnough, E. Johnson, Purification of Clostridium Botulinum type A neurotoxin, Methods in Molecular Biology (2000) 27–39.

[31] T. Grenda, A. Grenda, P. Krawczyk, K. Kwiatek, Botulinum toxin in cancer therapy—current perspectives and limitations, Applied Microbiology and Biotechnology 106 (2021) 485 – 495.

[32] Anti-clostridium Botulinum toxin A antibody [B364M] (ab252737), acessed on April 14, 2024 (2024).

[33] O. Rossetto, L. Morbiato, P. Caccin, M. Rigioni, C. Montecucco, Presynaptic enzymatic neurotoxins, Journal od Neurochemistry 97 (2006) 1534–1545.

[34] D. Dressler, R. Benecke, Pharmacology of therapeutic Botulinum toxin preparations, Disability and Rehabilitation 29 (2007) 1761–1768.

[35] J. Weisemann, N. Krez, U. Fiebig, S. Worbs, M. Skiba, T. Endermann, M. B. Dorner, T. Bergström, A. Munöz, I. Zegers, C. Müller, S. P. Jenkinson, M.-A. Avondet, L. Delbrassinne, S. Denayer, R. Zeleny, H. Schimmel, C. Åstot, B. G. Dorner, A. Rummel, Generation and characterization of six recombinant Botulinum neurotoxins as reference material to serve in an international proficiency test, Toxins 7 (12) (2015) 5035–5054.

[36] K. Czamara, K. Majzner, M. Z. Pacia, K. Kochan, A. Kaczor, M. Baranska, Raman spectroscopy of lipids: A review, Journal of Raman Spectroscopy 46 (1) (2015) 4–20.

[37] E. Wiercigroch, E. Szafraniec, K. Czamara, M. Z. Pacia, K. Majzner, K. Kochan, A. Kaczor, M. Baranska, K. Malek, Raman and infrared spectroscopy of carbohydrates: A review, Spectrochimica Acta Part A: Molecular and Biomolecular Spectroscopy 185 (2017) 317–335.

[38] J. Zhang, H. Jiang, P. Gao, Y. Wu, H. Sun, Y. Huang, X. Xu, Confocal Raman microspectroscopy combined with chemometrics as a discrimination method of clostridia and serotypes of Clostridium botulinum strains, Journal of Raman Spectroscopy 52 (11) (2021) 1820–1829.

[39] M. R. M. Khorasan, M. Rahbar, A. Z. Bialvaei, M. M. Gouya, F. Shahcheraghi, B. Eshrati, Prevalence, risk factors, and epidemiology of food-borne Botulism in iran, Journal of Epidemiology and Global Health 10 (2020) 288–292.

[40] M. Young, K. Young, Scientific review of the aesthetic uses of Botulinum toxin type A, Archives of craniofacial surgery 22 (2021) 1–10.

[41] Y. Al-Tamimi, K. F. Ilett, M. J. Paech, S. J. O’Halloran, P. E. Hartmann, Estimation of infant dose and exposure to pethidine and norpethidine via breast milk following patient-controlled epidural pethidine for analgesia post caesarean delivery, International Journal of Obstetric Anesthesia 20 (2011) 128–134.

[42] K. Wanat, Biological barriers, and the influence of protein binding on the passage of drugs across them, Molecular biology reports 47 (4) (2020) 3221–3231.

[43] F. J. Nice, A. C. Luo, Medications and breast-feeding: Current concepts, Journal of the American Pharmacists Association 52 (1) (2012) 86–94.

[44] T. Whitmore, N. Trengove, D. Graham, P. Hartmann, Analysis of insulin in human breast milk in mothers with Type 1 and Type 2 diabetes mellitus, International journal of endocrinology 2012 (2012) 1–9.

[45] D. Intiso, Therapeutic use of Botulinum toxin in neurorehabilitation, Journal of Toxicology (2012).

[46] M. E. Duff, R. E. Morton, Managing spasticity in children, Paediatrics and child health 17 (12) (2007) 463–466.

[47] Y. C. Wang, D. Burr, G. J. Korthals, H. Sugiyama, Acute toxicity of aminoglycoside antibiotics as an aid in detecting Botulism, Applied and Environmental Microbiology 48 (5) (1984) 951–955.

[48] C. E. Doneanu, M. Anderson, B. J. Williams, M. A. Lauber, A. Chakraborty, W. Chen, Enhanced detection of low-abundance host cell protein impurities in high-purity monoclonal antibodies down to 1 ppm using ion mobility Mass Spectrometry coupled with multidimensional Liquid Chromatography, Analytical Chemistry 87 (20) (2015) 10283–10291.

[49] E. Brauchle, K. Schenke-Layland, Raman spectroscopy in biomedicine – non-invasive in vitro analysis of cells and extracellular matrix components in tissues, Biotechnology Journal 8 (2012).

[50] R. Ullah, S. Khan, S. Javaid, H. Ali, M. Bilal, M. Saleem, Raman spectroscopy combined with a support vector machine for differentiating between feeding male and female infants mother’s milk, Biomedical Optics Express 9 (2) (2018) 844–851.

[51] N. Feliu, M. Hassan, E. Garcia Rico, D. Cui, W. Parak, R. Alvarez-Puebla, SERS quantification and characterization of proteins and other biomolecules, Langmuir 33 (38) (2017) 9711–9730.

[52] D. G. Kotsifaki, R. R. Singh, S. N. Chormaic, V. G. Truong, Asymmetric split-ring plasmonic nanostructures for the optical sensing of Escherichia coli, Biomed. Opt. Express 14 (9) (2023) 4875–4887.

[53] M. Hallett, Explanation of timing of botulinum neurotoxin effects, onset and duration, and clinical ways of influencing them, Toxicon 107 (2015) 64–67.

[54] A. M. K. Rød, N. Harkestad, F. K. Jellestad, R. Murison, Comparison of commercial elisa assays for quantification of corticosterone in serum, Nature scientific reports 7 (1) (2017) 1–5.

